# Harnessing GPT-4 for Automated Curation of E3-Substrate Relationships in the Ubiquitin-Proteasome System

**DOI:** 10.1101/2024.10.20.619305

**Authors:** Zhiqian Zhang, Stephen J. Elledge

## Abstract

The ubiquitin-proteasome system (UPS) is a complex regulatory network involving around 600 E3 ligases that collectively govern the stability of the human proteome by targeting thousands of proteins for degradation. Understanding this network requires integrating vast amounts of information on gene and protein interactions scattered across unstructured literature. Historically, manual curation has been the gold standard for transforming such data into structured databases, but this process is time-consuming, prone to error, and unable to keep up with the rapid growth of scientific publications. To address these limitations, we developed a scalable, cost-effective workflow using GPT-4, a large language model (LLM), to automate the curation of degradative E3-substrate relationships from the literature. By mining approximately two million PubMed papers, we identified 7,829 degradation-related abstracts and curated a structured database of 3,294 unique E3-substrate pairs using GPT-4, achieving an annotation accuracy rate approaching that of human experts. The resulting database of E3-substrate pairs offers valuable insights into the ubiquitin-proteasome system by highlighting understudied E3s and previously unknown UPS substrates in proteome-wide stability experiments. This automated approach represents substantial increase in productivity compared to manual curation and stands as the largest effort to date utilizing LLMs for the automated curation of protein-protein regulatory relationships. We further showed that our approach is generalizable to other enzyme-substrate families, such as deubiquitinases, kinases, and phosphatases. Overall, our study demonstrates the potential of LLMs as a scalable technology for large-scale curation of signalling relationships, substituting and complementing manual curation to accelerate biological research.

## Introduction

Cellular signalling networks operate as complex, dynamic control systems, shaped by millions of years of evolution to regulate diverse biological processes. These naturally evolved networks resemble integrated circuits, where each component functions like a circuit element, transmitting signals that guide cellular responses to internal and external stimuli. Over the past few decades, molecular biology has focused on dissecting these networks by reverse engineering, breaking them down into simple relationships such as “gene A activates gene B” or “protein X inhibits protein Y.” This approach has generated a vast body of detailed knowledge about individual signalling relationships between genes. However, integrating this extensive information into an integrated database remains challenging due to the vast volume of published literature lacking easily accessible structured data and standardized reporting.

Historically, manual curation was the gold standard for transforming unstructured published data into structured databases^1–9^. However, manual curation is time-consuming, error-prone, and struggles to keep pace with the rapid growth of scientific publications^10^. Variability in literature reporting and manual curation further complicates standardization and integration. The exponential growth of scientific literature increasingly challenges the efficiency and feasibility of this approach^11,12^. Recently, the development of Large Language Models (LLMs) presents a new avenue for addressing these challenges^13,14^. Notably, GPT-4, a transformer-based LLM developed by OpenAI, has shown potential for the automated curation of various biological data^15–18^. However, its potential for curating gene/protein signalling relationships remains largely unexplored. It remains unclear if the technical challenges can be overcome, such as the difficulty in handling specialized domain vocabulary and the precision required in interpreting scientific texts.

The complex cellular signalling network operates via distinct biochemical mechanisms, organized into families with large combinatorial complexities. For instance, there are about 600 ubiquitin ligases (E3s) for ubiquitination of a wide variety of substrates^19^. A common outcome of ubiquitination is proteasomal degradation^20^, where ∼600 E3s collectively regulate the stability of ∼20,000 proteins in the human genome. Significant advances have been made in our understanding of the ubiquitin-proteasome system. However, it remains unclear how many degradative E3-substrates are documented, in part because this information is dispersed across unstructured text in thousands of publications.

Here, we established an ultra-fast, scalable, cost-effective workflow for automated curation of degradative E3-substrate relationships from literature using GPT-4, with accuracy approaching to that of human experts. Our analysis mined about two million PubMed papers, identified 7,829 degradation-related abstracts through text-mining, and curated 3,294 unique E3-substrate pairs into structured databases using GPT-4. This process, once set up, can be completed within a day at a cost of under $200. While we focused on ubiquitin ligases as a proof-of-concept, this workflow is generalizable to other domains, such as deubiquitinases, kinases, and phosphatases. Altogether, our work demonstrates the potential of LLMs for large-scale, cost-effective, automated curation of signalling relationships across complex families.

## Results

To systematically curate known degradative E3-substrate relationships, we first examined how these relationships have been historically identified. Some regulatory systems have been explored through comprehensive atlas projects, such as the BioPlex3.0 profiling of protein-protein interactions^21^, or through massively multiplexed assays, like genome-scale perturb-seq for transcriptional regulation profiling^22^. In contrast, the ubiquitin-proteasome system has remained largely inaccessible to such high-throughput approaches and has not been extensively characterized by any atlas project. High-throughput assays for mapping E3-substrate pairs in a multiplexed manner have only recently been developed^23^. As a result, most known degradative E3-substrate pairs were discovered individually and are often explicitly mentioned in the “abstract” sections of publications. We therefore hypothesized that large language models (LLMs), such as GPT-4, could be employed to curate unstructured information from abstracts, converting it into a structured form where E3-substrate pairs are specifically and uniquely defined using official gene symbols.

Preliminary testing with the OpenAI GPT-4 interface on individual abstracts suggested that GPT-4 can process some unstructured abstracts and extract E3-substrate pairs in the form of official gene symbols (Figure 1A). With human oversight, we iteratively refined our prompting strategy through a series of prompt engineering steps to optimize GPT-4’s performance across a broader set of abstracts. This effort culminated in a two-step chain-of-thought prompting strategy^24^ (See Methods). First, GPT-4 was prompted to input an abstract and identify any E3-substrate pairs mentioned, outputting them in the specific format “E3:substrate”. Next, GPT-4 was instructed to convert various E3 and substrate names from the first step into their official gene symbols. The prompts were specifically designed to improve GPT-4’s responses by: (1) ensuring that a degradation relationship between the E3 and the substrate is explicitly stated; (2) specifying the output format for E3 pairs, including scenarios where multiple pairs are described or none are mentioned; (3) ensuring GPT-4 only relies on information from the abstract for curation; and (4) guiding GPT-4 to consider specificity-conferring adaptors as “E3s” in the case of multi-subunit E3 ligases such as Cullin-RING ligases and Anaphase-Promoting Complexes (See Methods).

**Figure 1.**
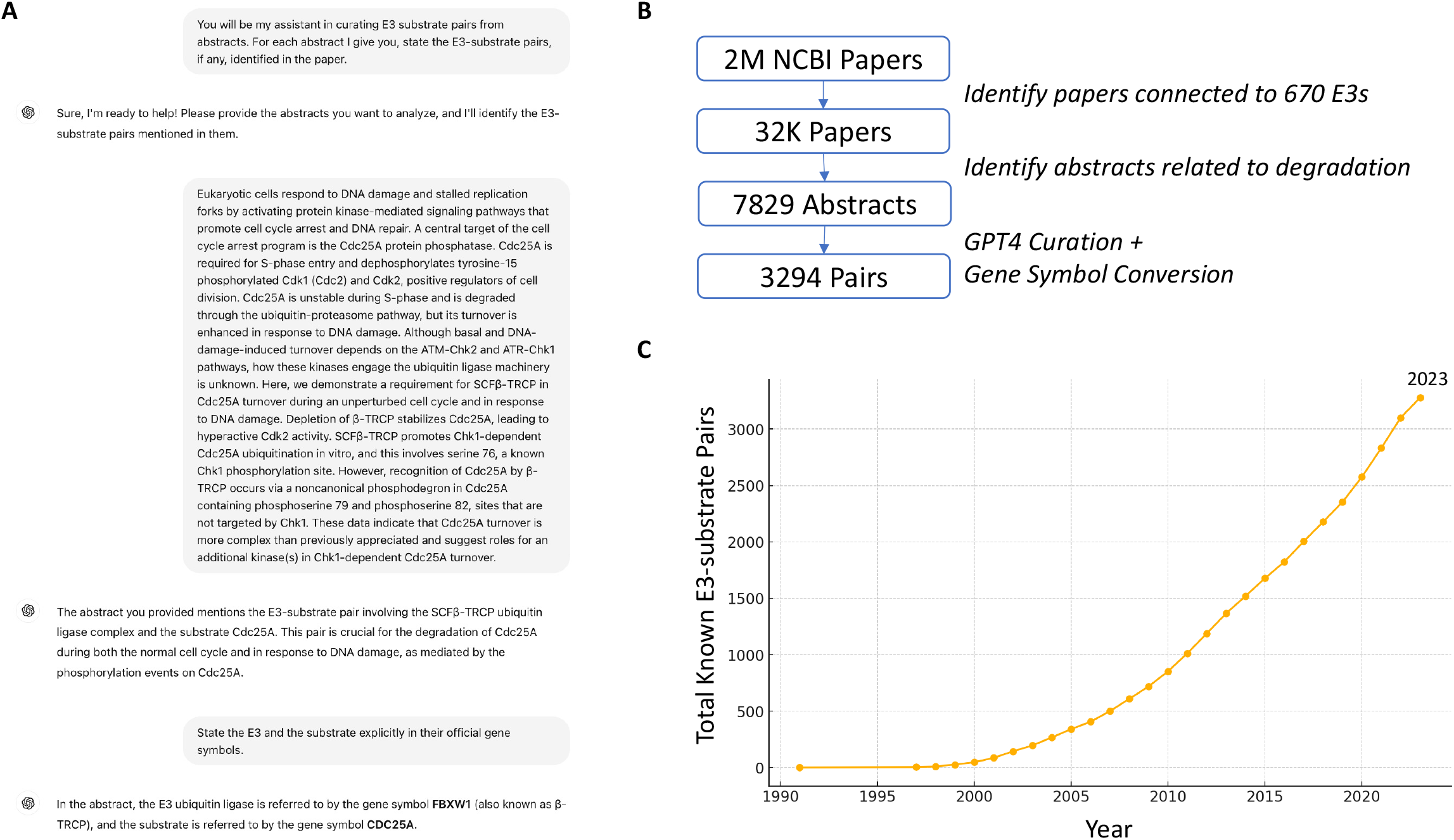
GPT-4 curated 3,294 degradative E3-substrate pairs from approximately 2 million PubMed articles. a) A visual capture of the GPT-4 user interface, demonstrating the model’s ability to curate E3-substrate pairs from abstracts. b) A schematic diagram illustrating the workflow for GPT-4-driven curation of degradative E3-substrate pairs. c) The cumulative number of known E3-substrate pairs from 1991 to 2024, as annotated in the GPT-4 curated database.

After demonstrating GPT-4’s capability for automated curation of individual abstracts, we aimed to create a comprehensive list of abstracts describing degradative E3-substrate relationships. The PubMed database contains approximately two million published papers related to human genes (See Methods). Theoretically, GPT-4 could read through all ∼2M abstracts and curate the degradative E3-substrate pairs, if any, from each abstract. However, the processing time and computational cost at this scale are substantial. To address this, we developed a two-step computational approach to generate a list enriched for papers characterizing individual E3-substrate pairs (Figure 1B). First, we extracted PMIDs annotated to be related to a previously curated list of 670 ubiquitin ligases in the human genome^19^ (Table S1A). Then, we retrieved the abstracts for these PMIDs and filtered for those containing keywords related to protein degradation (i.e., degrad|destab), resulting in 7,829 PMIDs (Table S1B).

The GPT-4 user interface is not designed for large-scale prompting interactions. To scale up the curation to process the 7,829 abstracts, we utilized the GPT-4 API, sending each abstract to the OpenAI server for GPT-4 curation of E3-substrate pairs individually (Table S1B). The resulting thousands of names of E3s and substrates were sent back to the OpenAI server in batches for conversion into their corresponding “official gene symbols” by GPT-4 (Figure 1B; Table S1C). Since GPT-4’s conversion of names into gene symbols was not always accurate (e.g., different aliases were sometimes used for the same gene), the “official gene symbols” curated by GPT-4 were first converted to official gene IDs, and were then converted back to official gene symbols to ensure a list of accurately defined E3-substrate pairs where each gene is represented by a single, consistent official gene symbol, without aliases (See Methods). The resulting E3-substrate pairs were further filtered using the initial list of 670 E3s to retain only those pairs in which the E3 is among the 670 specified E3s. This yielded a preliminary list of 3,294 unique E3-substrate pairs described in 7,829 degradation-related abstracts (Figure 1B), which is comprehensively summarized in Table S1D in a searchable fashion. The number of discovered E3-substrate pairs increased exponentially since 2000s (Figure 1C).

To assess the accuracy of the systematic curation, we randomly sampled 100 curated E3-substrate pairs and manually assessed the accuracy of curation (Figure S1). A curation is considered as accurate if the abstract implies that the E3 degrades or destabilizes the substrate. Remarkably, 83% (83 out of 100) were found to be accurate (Figure 2A). Errors can be broadly classified into three types that closely resemble human errors: (1) Gene symbol conversion errors—GPT-4 correctly identified the regulatory relationships but returned incorrect official gene symbols; (2) Misunderstandings—GPT-4 misinterpreted the types of regulatory relationships or the proteins involved as described in the abstracts; (3) Over-interpretations—the abstracts did not clearly specify the types of regulatory relationships or the proteins involved, yet GPT-4 over-interpreted the information. (Figure 2A; Figure S1; Table S1E). Altogether, these findings suggest that the accuracy of GPT-4’s curation is approaching that of a human expert. However, more advanced prompt engineering will be required to guide GPT-4 through the nuances of gene nomenclature and convoluted abstracts..

**Figure 2.**
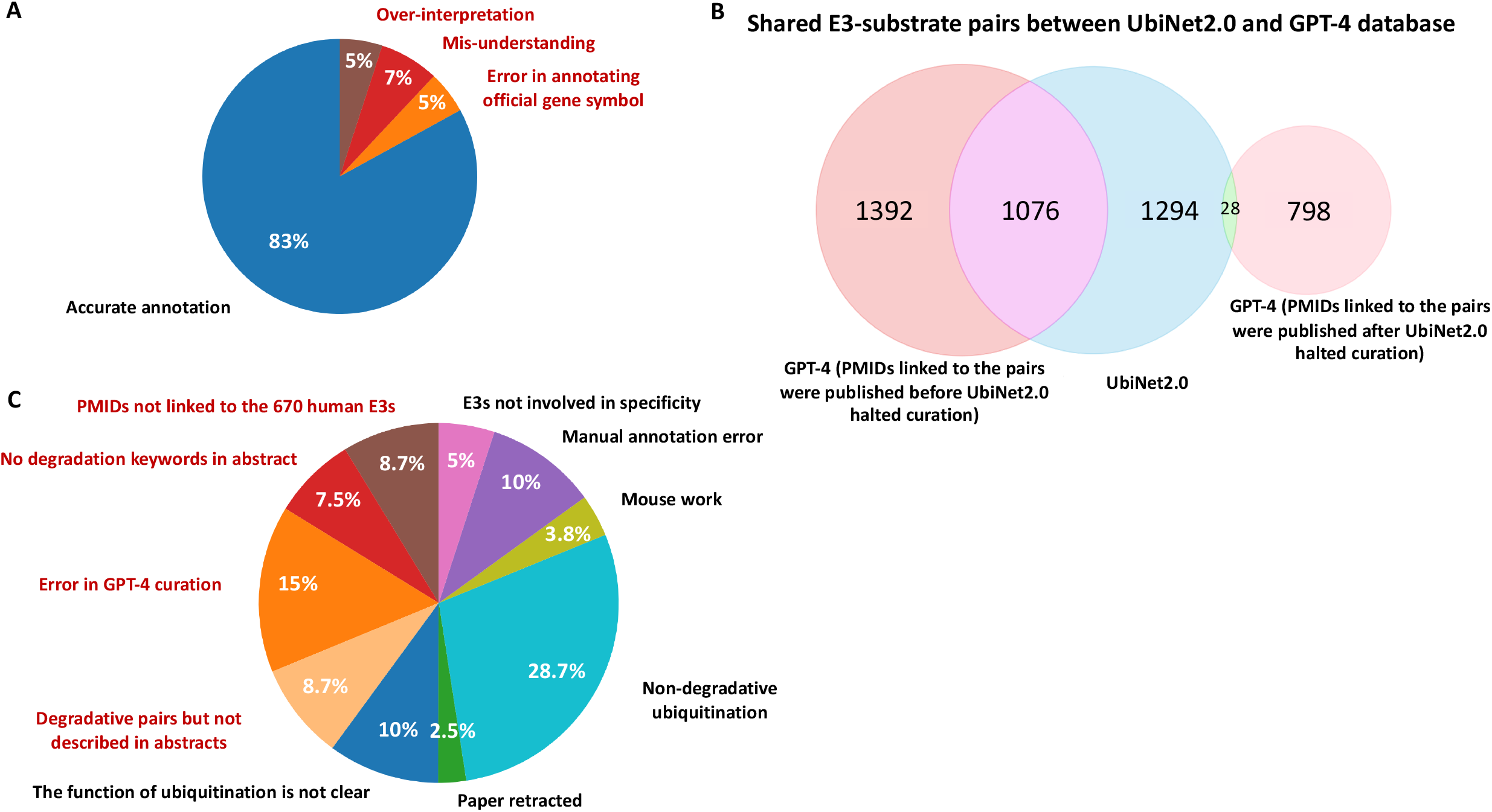
The accuracy of GPT-4 curation closely approximates that of human experts. a) Out of 100 randomly sampled GPT-4 curated pairs, 83 were manually verified as accurate. Different underlying reasons resulting in the inaccurate annotation were annotated in red. b) A comparison between the GPT-4 curated E3-substrate database and the manually curated UbiNet2.0 E3-substrate database. UbiNet2.0 halted curation in June 2020 and did not differentiate between degradative and non-degradative pairs. c) Out of 80 randomly sampled pairs curated in UbiNet2.0 but missed in GPT-4 curation, 40% were false negatives (true degradative E3-substrate pairs), 60% were true negatives (they should not be considered as degradative E3-substrate pairs). Different types of main underlying reasons for the inaccurate classifications (false negatives, colored in red, or true negatives, colored in black) were annotated.

We compared the GPT-4-curated database with UbiNet2.0^25^, the most comprehensive and up-to-date E3-substrate database, which manually curated 2,409 E3-substrate interactions in humans involving 367 E3s (Table S1F), but did not differentiate between degradative and non-degradative ubiquitination. Up until June 2020, when UbiNet2.0 ceased its curation efforts, the GPT-4 curation identified 1,076 pairs that were also annotated in the UbiNet2.0 database (Figure 2B). Notably, GPT-4 curation identified an additional 1,392 pairs documented before June 2020 that were missed by the manual curation in UbiNet2.0. This discrepancy is likely due to GPT-4’s use of more inclusive keywords (e.g., terms containing “degrad” or “destab”) in its literature search, compared to the more restrictive keyword criteria used in UbiNet2.0^25^, which were likely designed to improve the relevance of papers that were subjected to manual curation. Furthermore, GPT-4 has identified 798 additional pairs that have been discovered since UbiNet2.0 stopped curating (Figure 2B). Interestingly, UbiNet2.0 included 1,294 pairs that were not captured by the GPT-4 curation (Figure 2B). These pairs can be classified into false negatives (degradative pairs missed by GPT-4 curation) and true negatives (pairs that should not be considered conclusively degradative based on the linked paper). To estimate the percentage of false negatives and understand why they were missed, we randomly sampled 80 pairs that were annotated in UbiNet2.0 but not in the GPT-4 database. Among these, 32 were identified as false negatives, while 48 were true negatives (Figure 2C; Figure S2; Table S1G). The true negatives included E3-substrate pairs that were non-degradative regulations or functionally unclear according to the linked papers and manual annotation errors. Additionally, some degradative relationships were exclusively studied in mouse or involved E3s not conferring substrate specificity such as ARIH2 and RBX2, and were therefore not annotated in the GPT-4 database as it was originally designed. The false negatives (Figure S2) can be categorized into four main types: (1) Errors in GPT-4 curation, where GPT-4 processed the abstracts of the linked PMIDs but failed to annotate the correct pair, which were often due to errors in converting various gene names to their official gene symbols; (2) Abstracts without pre-determined degradation keywords and thus were not sent to GPT-4 for downstream curation; (3) PMIDs not linked to the 670 human E3s—cases where the abstracts described E3-substrate pairs but were not associated with the 670 human E3s in the PubMed database; and (4) Degradative E3-substrate pairs that were not described in the abstract. Overall, these findings suggest that while GPT-4-based automated curation can systematically rediscover a great proportion of manually curated E3-substrate pairs and identify numerous additional pairs missed by manual efforts or due to outdated databases, further optimizations are needed to reduce the number of false negatives – degradative E3-substrate pairs missed by the current approach.

We reasoned that this curated database of degradative E3-substrate pairs provides a rich dataset for systematic analysis of UPS research over the years (Figure 3A). We first analysed the number of known substrates for each E3. The E3s with the most known substrates include STUB1, BTRC, MDM2, FBXW7, NEDD4, PRKN, CBL, SKP2, VHL, FZR1, SYVN1, ITCH, SMURF1, NEDD4L, TRIM21, SIAH1, SPOP, SIAH2, SMURF2, and UBE3A, each with more than 40 known substrates (Table S2A). Plotting the number of known substrates for each of the 670 E3s revealed a long-tail distribution (Figure S3A): while 60 E3s have more than 10 known substrates and 62 E3s have 6-10 substrates, 272 have only 1-5 substrates (Figure 3B). Strikingly, 276 E3s have no known substrates at all (Figure 3B). This includes many members of the Kelch, MAGE, RNF, and TRIM families (Table S2A). DAVID analyses^26^ of the substrate lists for some of the extensively studied E3s revealed significantly functional enrichment consistent with the known functions of the E3s (Figure 3C), suggesting the potential of systematic functional enrichment analyses in generating functional hypotheses for each E3s using the curated substrate lists. We analysed the number of known E3s for each substrate and found a long-tail distribution (Figure S3B). Notably, substrates such as TP53, CTNNB1, MYC, CDKN1A, and EGFR each have over 20 associated E3s, a biologically unlikely scenario that suggests some of the curated E3-substrate pairs may not result from direct interactions. Further refinements are needed to distinguish pairs with biochemical evidence of direct interactions from those without. A positive correlation was observed between the number of known E3s and the number of PMIDs associated with each substrate (Figure S4: Table S2B), indicating that extensively studied substrates tend to have more identified E3s. Additional E3-substrate pairs will likely be discovered once understudied proteins are further investigated.

**Figure 3.**
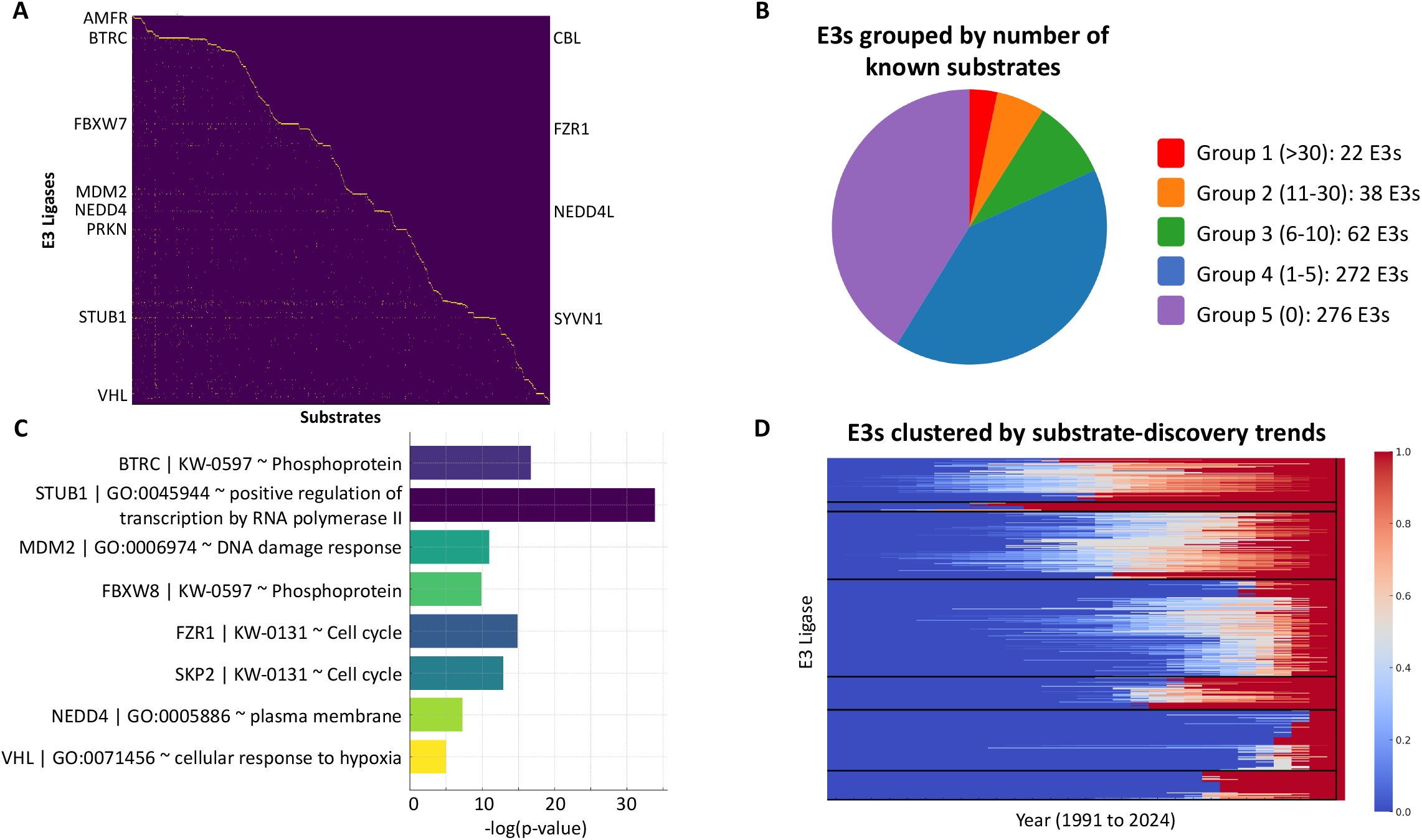
Analyses of the E3s using the list of GPT-4 curated E3-substrate pairs. a) E3-substrate pairs curated by GPT-4 are shown. Representative E3s with many known substrates were annotated. b) E3s categorized by their number of known substrates. c) Analysing the substrate lists for each selected E3s using DAVID identified significantly enriched terms consistent with the known biological functions of the E3s. For each term enrichment, the -log(p-value) is shown. d) A temporal analysis of substrate discovery trends for each E3, clustering them into distinct groups. The heatmap represents the cumulative number of known substrates of E3 ligases over the years, normalized to their total number in 2024 (i.e., Year 2024 = 1 for each E3s). E3s were classified into seven distinct clusters based on their temporal substrate discovery patterns using k-means clustering.

The total number of known E3-substrate pairs has increased exponentially since the initial discovery in 1991^27^ (Figure 1C). We reasoned that E3s can be clustered into distinct groups based on when their first substrate was discovered and the rate of substrate discovery since then. Therefore, we counted the total number of known substrates for each E3 each year, normalized them to the total number of known substrates for each E3 up to 2024, and clustered the E3s based on their substrate discovery trends over time (Figure 3D: Table S2C). This analysis revealed multiple E3 groups with distinct substrate-discovery trends (Figure 3D;; Table S2D).

The curated database could be integrated with systematic protein stability assays^28–31^ to generate a prioritized list of unstable proteins without known cognate E3s. As a proof of concept, we analysed 994 unstable proteins identified by deep quantitative proteomics^28^ and found that 204 of these proteins have known E3s (Figure 4; Table S2E). Of the remainder, 89 proteins are ubiquitin ligases themselves and are likely unstable due to self-ubiquitination. Remarkably, 701 proteins neither have known E3s nor are ubiquitin ligases, highlighting them as attractive candidates for downstream biochemical and genetic analyses to identify their cognate E3s.

**Figure 4.**
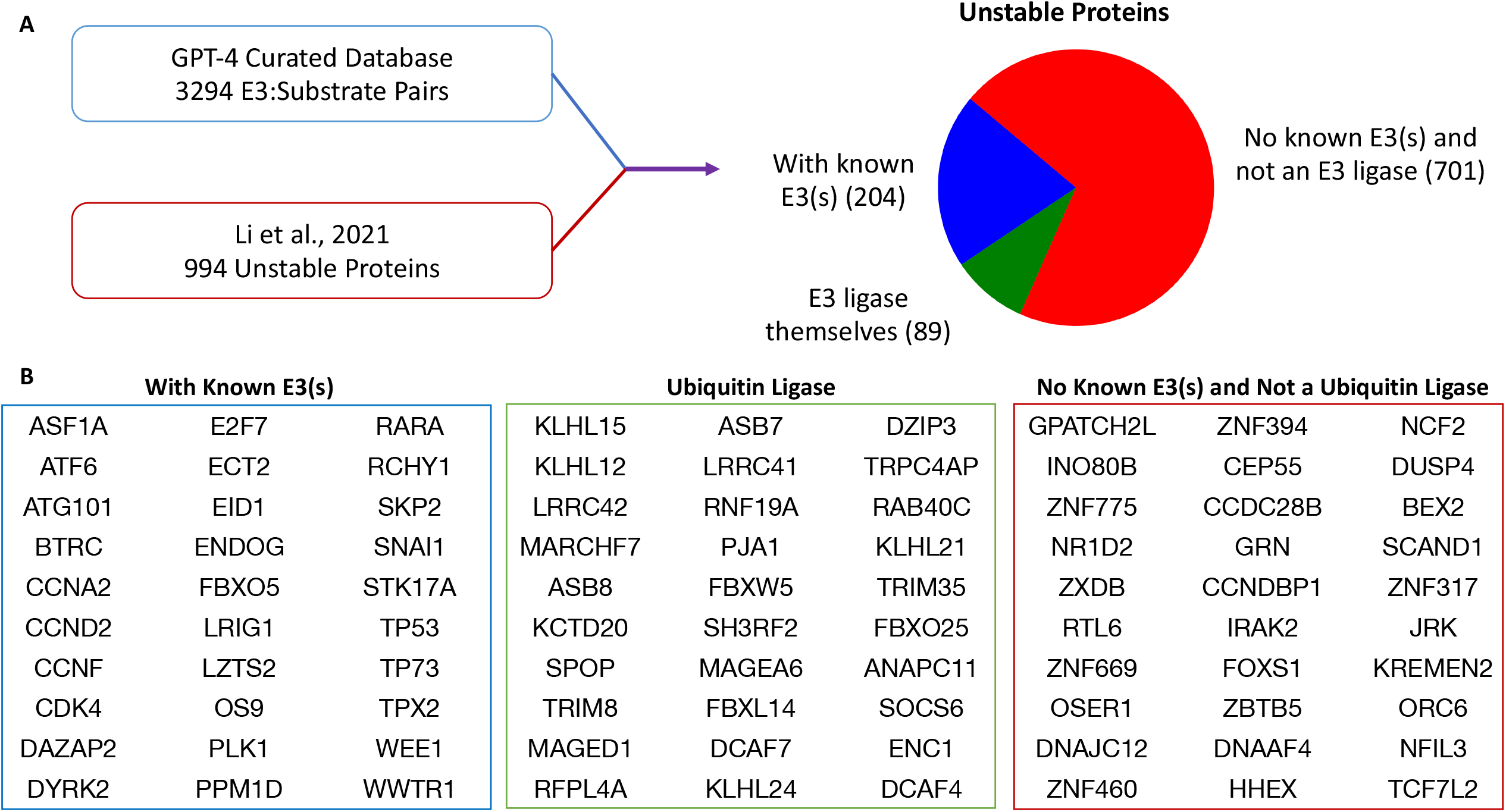
Application of the GPT-4-curated database to a proteome-wide stability dataset identifies known UPS substrates and novel candidate substrates. a) Integration of the GPT-4-curated database with 994 unstable proteins from Li et al., 2021^28^ identified 204 unstable proteins with known E3s, 89 unstable proteins that are ubiquitin ligases, and 701 unstable proteins that neither have known E3s nor are ubiquitin ligases. b) Thirty randomly sampled proteins are presented for each group described above.

Although this study focused on curating degradative E3-substrate pairs, we speculated the same analytical framework can be generalized to curate any complex system that has traditionally relied on manual curation of massive unstructured text data. The human genome encodes about a 100 deubiquitinases^32^, 500 kinases^33^, and 200 phosphatases^34^, each regulating a wide variety of substrates. While curating databases for each of these families is beyond the scope of this paper, we demonstrated that GPT-4 can curate specific enzyme-substrate pairs for all three families using one representative abstract from each as an example (Figure S5).

## Discussion

The signalling relationships between proteins and the challenges in characterizing them represent one of the most complex areas of modern research. Over the past 30 years, increased biological research activities, combined with decreasing data storage costs and the rise of the internet, has led to an exponential growth of openly accessible biological data on protein-protein regulatory relationships. High-throughput technologies, such as quantitative proteomics and high-throughput screening platforms, systematically profile these relationships and generate large-scale omics datasets. However, the full potential of such datasets cannot be realized without the ability to easily distinguish novel relationships from known ones. While manual curation of published papers has proven effective for guiding future research, it has significant drawbacks: it is time-consuming, labour-intensive, and prone to human error. Moreover, such manual curation efforts cannot be easily adapted for personalized downstream analyses, a feature that is increasingly in demand.

Despite the extensive efforts to manually curate hundreds or thousands of E3-substrate relationships in databases such as UbiNet2.0^25^ and UbiBrowser^35^, these efforts did not distinguish degradative from non-degradative relationships, and replicating such extensive curation can be time-consuming and expensive. Here, using degradative E3-substrate relationships as a proof of concept, we present an ultra-fast, automated, scalable, and cost-effective approach for curating known E3-substrate pairs from published abstracts. Once established, this analysis can be completed automatically within 1 day at a cost of less than $200, thus offering a substantial productivity boost of both time and cost and democratizing large-scale curation by making it accessible to any researcher within a single day and at minimal expense.

The automated curation pipeline presented here achieved an annotation accuracy approaching that of human experts (∼83%). Optimization strategies such as chain-of-thought prompting^24^ and specific instructional prompts were used to enhance annotation accuracy. Most remaining inaccuracies arose from errors in gene symbol conversion for specific E3s and substrates, or occasional mis-understandings and over-interpretations of the abstracts. We anticipate that ongoing improvements in GPT models and other LLMs, coupled with more refined prompt engineering involving few-shot learning^36,37^, chain-of-thought prompting^24^ and self-consistency^38^, will further enhance annotation accuracy to nearly 100%. Comparison with the manually curated UbiNet2.0 database revealed that certain degradative E3-substrate pairs were missed by the GPT-4 curation pipeline. One of the main causes of these “false negatives” is curation errors, where GPT-4 failed to process the information contained in the abstracts into E3-substrates specified by official gene symbols. This could likely be reduced with more refined prompt engineering techniques used to improve curation accuracy. Another contributing factor is that papers describing degradative pairs but not linked to the list of 670 E3s in the PubMed database were excluded. This issue could potentially be addressed by using a more inclusive approach for abstract collection for downstream GPT-4 processing. Lastly, although uncommon, degradative E3-substrate pairs mentioned only in the main text were inevitably missed by the current pipeline.

In addition to annotation errors and false negatives, multiple limitations are present with our current approach. Focusing solely on the abstract section inherently limits this curation pipeline by excluding potentially informative details found in the main text, such as the quantitative degree of degradation, model organisms and cell types used in degradation studies, the degradation pathway involved (proteasomal or autophagy), conditions required for active degradation and others. The presented approach also did not assess the presence of experimental evidence supporting a direct E3-substrate relationships such as biochemical data on E3-substrate interactions and substrate ubiquitination. These limitations must be considered, and the curated E3-substrate pairs should be regarded as “candidate” pairs. In the near future, we expect that advanced curation pipelines using multimodal AI models such as GPT-4o or models that emphasize chain-of-thought reasoning such as GPT-o1, could enable sophisticated curation that interprets image-format figure data and entire manuscript texts with strong reasoning capabilities.

The 3,294 degradative E3-substrate pairs curated here offer several immediate benefits to the UPS research community. First, they allow for intriguing observations from a system biology perspective. For example, why do some E3s have tens or hundreds of known substrates while many others have fewer than five? Do some E3s naturally have more substrates, or is this a reflection of the research focus and history of UPS research? It will be informative to apply high-throughput technologies to systematically define the substrates of ubiquitin ligases with relatively few known substrates^30,39–42^. Many extensively studied E3s, such as those involved in cell cycle progression and growth signalling (e.g., BTRC, MDM2, FBXW7, SKP2, FZR1, SPOP), were likely highlighted partly due to intensive research in cell growth and cancer biology^43^. Additionally, how many more E3-substrate pairs remain to be discovered? Comparing the curated database with other omics datasets profiling degradation ^28–31^ can also accelerate the discovery of novel E3-substrate pairs. For example, we showed that cross-referencing the database with an omics dataset that profiles 994 unstable proteins using deep quantitative proteomics^28^ immediately identifies many unstable proteins without known cognate E3s. These proteins can then be targeted for downstream genetic and biochemical analyses to map their E3s. Comparing this curated database with systematic degron datasets would also be valuable in identifying how many known E3 substrates have known degrons^44–47^.

Although LLMs have been applied to automated curation in biology, to our knowledge, none have achieved the scale demonstrated in this study—curating 3,294 regulatory gene pairs of the same family from 7,829 abstracts, with accuracy approaching that of human experts. During the preparation of this manuscript, we identified a conceptually similar preprint that used GPT-4 to curate the Medical Action Ontology for rare diseases from 4,918 medical abstracts^56^. Together, these studies highlight the potential for LLM-powered automated curation to be expanded to other complex areas of biological research that have traditionally relied on labour-intensive manual curation^1–9^. With advancements in accuracy and computational power, we anticipate that automated curation will both complement and eventually replace many manual efforts, relieving biologists from the repetitive task of searching and organizing information. This paradigm shift will provide more time for creative scientific exploration driven by human insight, thereby accelerating biological and biomedical research to the next level.

## Supporting information

GPT-4 curation of candidate E3-substrate pairs.

Analyses of curated E3-substrate pairs.

## Supplementary Figures

**Figure S1.**
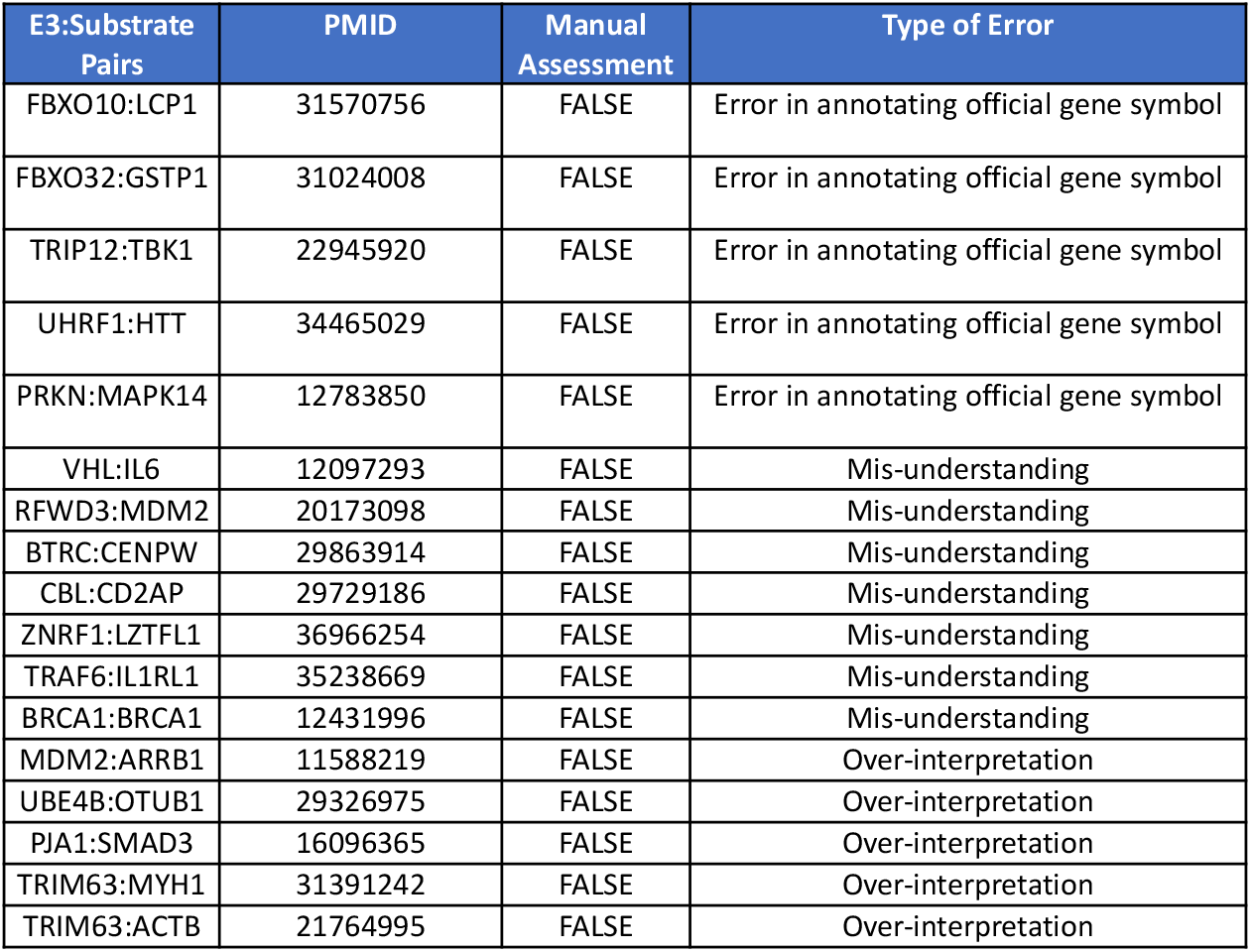
Manual assessment of the GPT-4 curation accuracy using 100 randomly sampled pairs. Different types of curation errors are shown.

**Figure S2.**
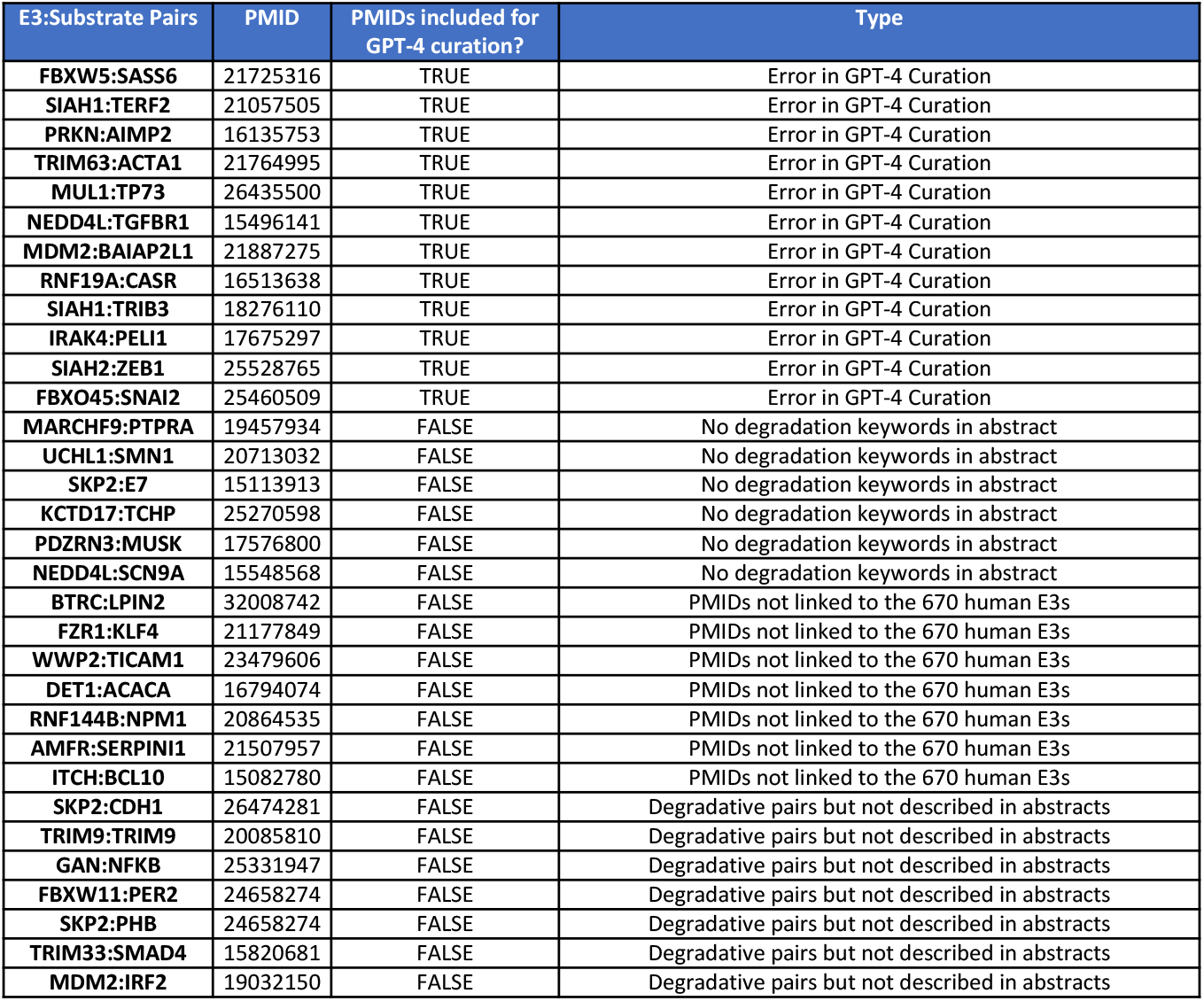
Manual assessment of pairs curated in UbiNet2.0 data but missed by GPT-4 curation using 80 randomly sampled pairs. False negatives due to different types of errors are shown.

**Figure S3.**
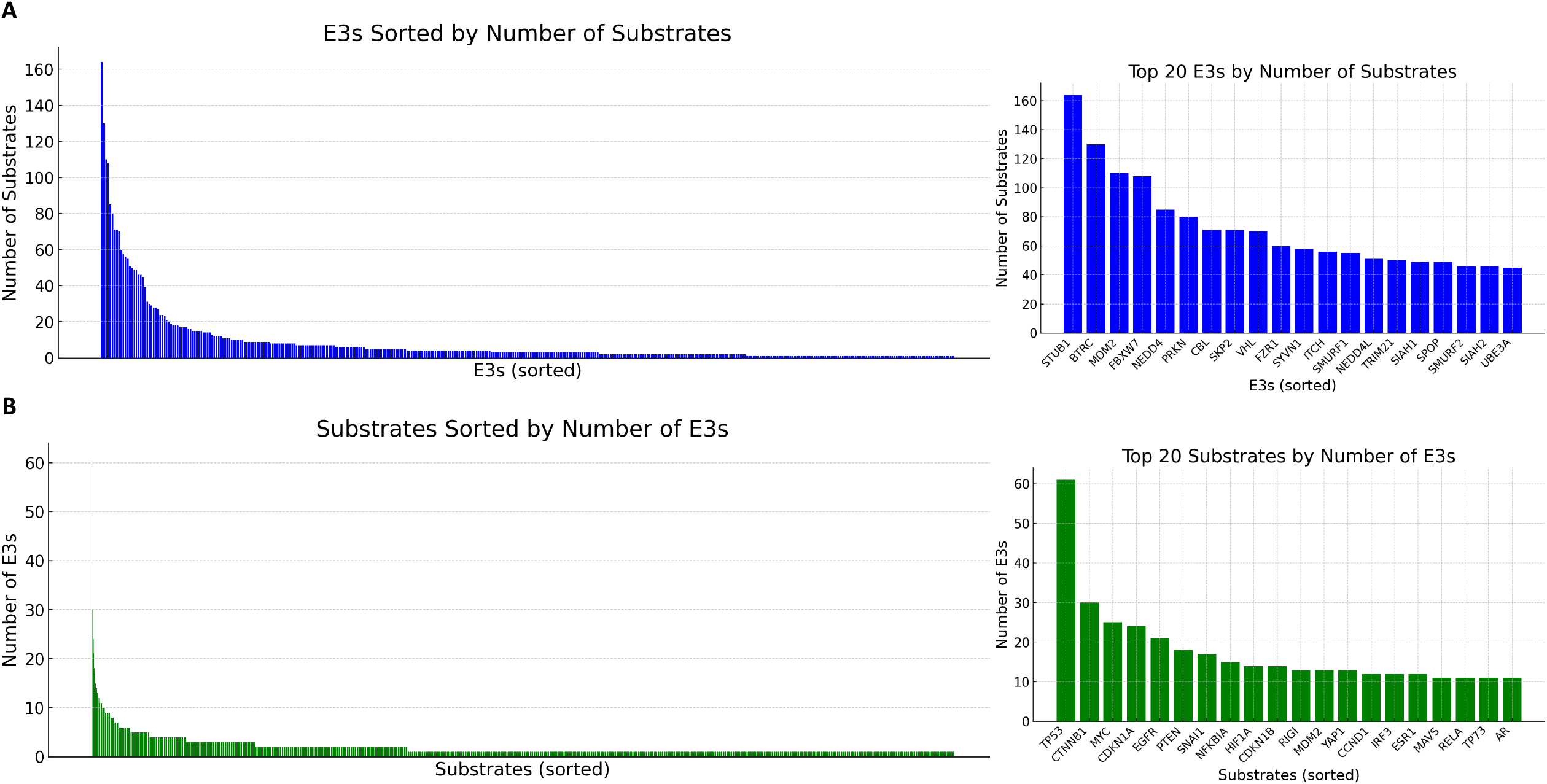
Plots showing (A) the number of substrates for each E3s and (B) the number of E3s for each substrates.

**Figure S4.**
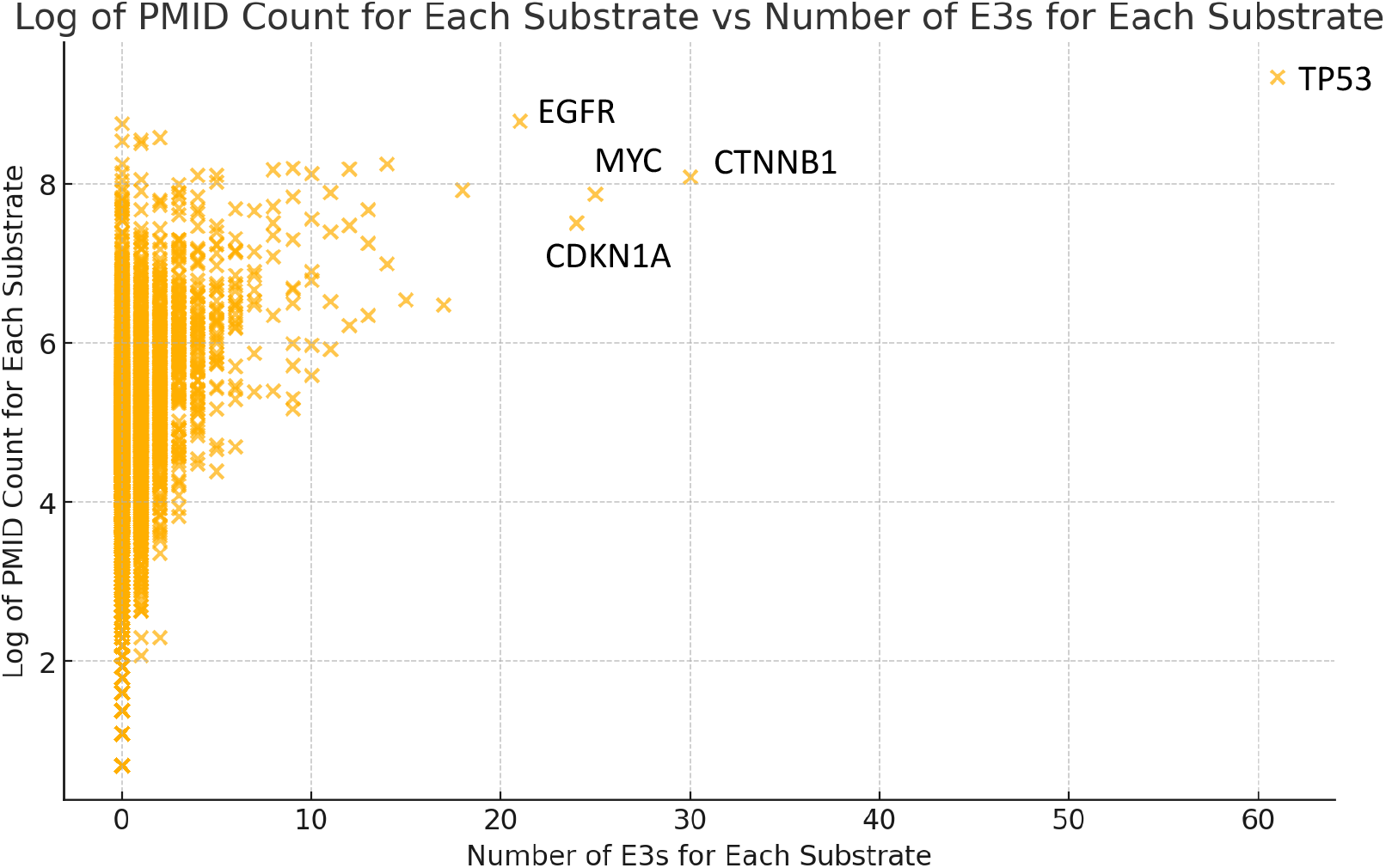
Plot showing the correlation between the number of E3s and the number of PMIDs for each human protein. Each dot represents a human protein coding gene.

**Figure S5.**
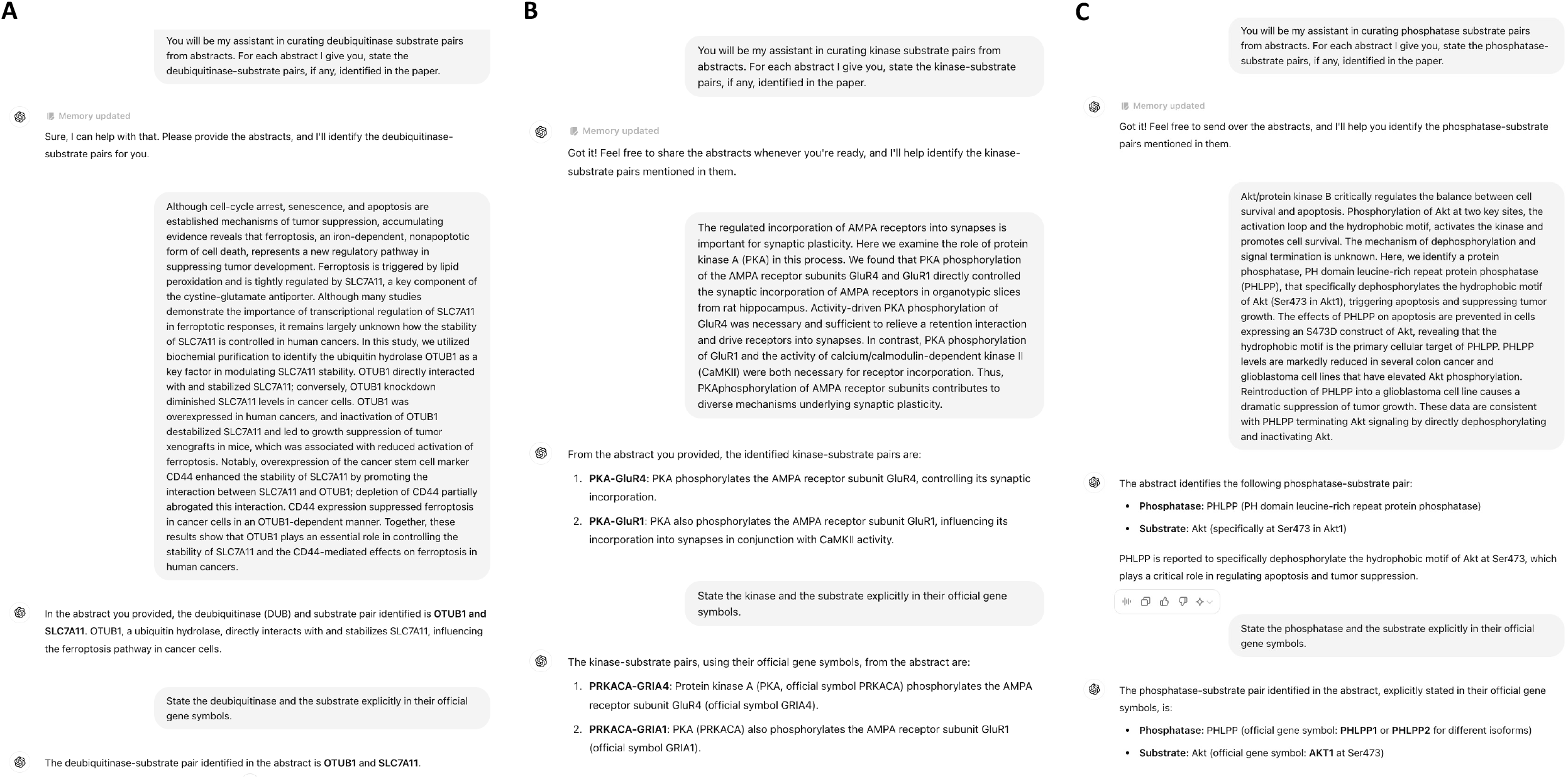
Preliminary testing suggested that GPT-4 can be applied to curate enzyme-substrate pairs for (A) deubiquitinases, (B) kinases and (C) phosphatases.

## Supplementary Tables

**Table S1. GPT-4 curation of candidate E3-substrate pairs**.

a) List of 670 E3s in gene IDs.

b) Preliminary GPT-4 curation results.

c) Conversion scheme from gene names to gene symbols using GPT-4.

d) Final list of 3,294 candidate E3-substrate pairs.

e) Manual assessment of 100 E3-substrate pairs randomly sampled from the 3,294 curated pairs

f) Comparison between UbiNet2.0 and the GPT-4 curated database.

g) Manual assessment of 80 randomly sampled E3-substrate pairs annotated in UbiNet2.0 but not in the GPT-4 database.

**Table S2: Analyses of curated E3-substrate pairs**.

a) Total number of known substrates for each E3s.

b) Total number of known E3s for each protein.

c) Normalized substrate discovery trends for each E3s.

d) Classification of E3s based on their substrate discovery trends.

e) Cross comparison with quantitative proteomics profiling proteome half-lives.

## Methods

### Collection of degradation-related abstracts

A list of 2,044,430 PMIDs linked to human genes (Organism ID: 9606) were retrieved from PubMed on Feb 7, 2024. The UbiHub collection of 670 E3s in human genome was retrieved from Liu et al. (2023)^19^. A list of 57,671 PMIDs linked to the 670 E3s were identified. Corresponding abstracts were retrieved from PubMed using Biopython Entrez utilities and XML records were parsed to extract text^57^. Text mining identified a list of 7,829 abstracts containing the keywords “degrad” or “destab”.

### Curation of E3-substrate pairs from abstracts using GPT-4

Each abstract was first sent to GPT4-turbo with the following prompt: [user_request = “You will be my assistant in curating E3 substrate pairs from abstracts. The definition of a E3 substrate pair in this context is that the E3 ubiquitinates and degrades the substrate, and make sure the degradation relationship is explicitly stated in the abstract. If you could curate multiple E3:substrate pairs from the same abstract, state the answer as E3_A:Substrate_A, E3_B:Substrate_B, etc… Your answer should only have one “:” per pair. Return NA if you cannot conclude. Do not use external knowledge other than the given abstract.”].

The collection of gene names curated by GPT4-turbo was converted to official gene symbols using GPT4 by the following prompt: [user_request = “I want you to be my assistant in gene symbol conversion. I will send you a list of gene names or a collection of gene names separated by ,. Substitute each collection with one official gene symbols in human. Listen carefully for the guidelines. For a collection of gene names, ignore terms including F-box, SCF, CUL1, CUL2, CUL3, CUL4A, CUL4B, CUL5, CRL, RBX1, RBX2, APC, APC/C, DDB1, ElonginB/C, EloBC, look for other E3 ligases in the entry connected by - or bracket. Pay special attention to gene names inside brackets for conversion. Always convert to ONE gene symbol by picking your best guess and state NA if you cannot. Always choose the most commonly associated official HGNC symbol. Your response should be in the format Gene Name A; Gene Symbol A, Gene Name B; Gene Symbol B.”]

The GPT4 ignored the conversion for certain non-typical gene names, and those were sent back to GPT4 iteratively for conversion until all gene names were converted using the following prompt: [user_request = “I want you to be my assistant in gene symbol conversion. I will send you a list of gene names or a collection of gene names separated by ,. Substitute each collection with one official gene symbols in human. Listen carefully for the guidelines. For a collection of gene names, ignore terms including F-box, SCF, CUL1, CUL2, CUL3, CUL4A, CUL4B, CUL5, CRL, RBX1, RBX2, APC, APC/C, DDB1, ElonginB/C, EloBC, look for other E3 ligases in the entry connected by - or bracket. If you see APC or APC/C, focus on if you see CDH1, CDC20, FZR1 nearby. If not, simply convert to APC/C. Pay special attention to gene names inside brackets for conversion. Always convert to ONE gene symbol by picking your best guess and state NA if you cannot. Always choose the most commonly associated official HGNC symbol. Do not skip any conversion task. Your response should be in the format Gene Name A; Gene Symbol A, Gene Name B; Gene Symbol B.”].

Gene symbols curated by GPT-4 were converted to gene IDs using the PubChem Gene Symbol to Gene ID Conversion Tool^58^. Before the conversion, “CDH1” was first manually converted to “FZR1” because of the confusing nomenclature between Cadherin-1 (CDH1) and CDC20 homolog 1 (FZR1). Afterwards, gene IDs were converted to gene symbols using the SynGO portal^59^ to generate a list of E3-substrate pairs with consistent official gene symbols. This list were then filtered to remove any “E3-substrate” pairs which the “E3s” are not part of the starting list of 670 E3s. This filtered list contained the final list of 3,294 curated E3-substrate pairs in their official gene symbols.

### Manual assessment of GPT-4 curation accuracy

A total of 100 curated E3-substrate pairs were randomly sampled from the list of 3,294 curated pairs and manually assessed for the accuracy of curation. For pairs with multiple PMIDs, the first PMID was used to assess the curation accuracy. A curation is considered as accurate if the abstract implies that the E3 degrades or destabilizes the substrate (Figure S1). Candidate E3-substrate pairs requiring mutants were noted but considered as “accurate” annotation, as they were considered as the broader group of “conditions required for degradation”. A total of 80 E3-substrate pairs were randomly sampled from the list of 1,294 pairs curated in UbiNet2.0 but not in the GPT-4 database. Pairs linked to PMID:22199232 were not included as that was the E3Net database paper the UbiNet2.0 built upon. The 80 pairs linked to 80 abstracts were manually assessed and classified into different types of reasons causing them to be missed by the GPT-4 curation (Table S1G).

### Temporal analysis of E3s substrate discovery trends

Multiple PMIDs linked to the same E3-substrate pairs were grouped together. The discovery year for each E3-substrate pair was defined as the year of the earliest linked publication. For each E3s, the total number of known substrates for each year were first counted, and normalized to their total number of known substrates in 2024. The E3 ligases were then clustered into 7 groups using k-means clustering and manually analysed to identify pattern in the data over the years.

## Author Contributions

Z.Z.. and S.J.E. conceptualized the project. Z.Z. performed all the computational analyses. Z.Z.. and S.J.E. wrote the manuscript. S.J.E. supervised the study.

## Acknowledgments

Z.Z. is a Croucher Ph.D. scholar. S.J.E. is a member of the Ludwig Center at Harvard and an Investigator with the Howard Hughes Medical Institute.

